# Modelling benefits and costs of decision making and feedback control for organisms in changing environments

**DOI:** 10.1101/2024.11.30.626137

**Authors:** Rajneesh Kumar, Iain G. Johnston

## Abstract

Cells (and organisms) make decisions in response to their environments. These decisions may help organism survival in environments with limited resource, but also constitute a cost to the organism in terms of the energy involved in sensing, processing, and responding to environmental change. Here, we explore the tradeoffs involved in a cost-benefit analysis of model organisms facing challenging deterministic or stochastic environments. The benefits of tunable versions of proportional-integral-derivative (PID) control are computed under different environmental behaviours; the model reflects both the ability to use this control to decide on cellular strategy and the potential cost associated with this feedback control. We quantify the circumstances under which control is most and least beneficial and the different weightings of the PID terms that perform best in specific and general situations. While our model is very simple, these results provide potential insight into the benefits of different control mechanisms, particularly at the single-cell level.

## Introduction

The ability of populations to adapt and respond to environmental change is central to ecology and evolution. Biological examples abound, across length and time scales, of organisms that change their level of activity in response to challenging environments. These examples include “persister cells” in bacteria, which adopt a less active state in an attempt to survive environmental challenges (Balaban et al., 2004; Veening et al., 2008), hibernation in animals, where a long dormancy period is adopted during predicted times of reduced resource availability (Boyles et al., 2020; Geiser, 2013); to extremes like tardigrade dormancy, which can last years without loss of organismal viability (Guidetti et al., 2011). Challenges of limited environmental resource can spark fierce competition (Chesson & Huntly, 1997), the evolution of efficient resource utilization (Schoener, 2011) and robust energetic machinery (García-Pascual et al., 2022), and the ability to adapt to different environments (Svanbäck & Bolnick, 2007). One particular adaptive strategy in the face of environmental change is bet-hedging (Cohen, 1966; Slatkin, 1974), where cells (or organisms) adopt different phenotypes or behaviours, increasing the chance that some will survive in an unpredictable future (Ackermann, 2015; Beaumont et al., 2009).

Sensing, predicting, and adapting to different environments involves processing information, which requires energy (Landauer, 1961; ten Wolde et al., 2016; Wolpert et al., 2024), and responding to this processing by making cellular decisions, which also requires energy (Forrest et al., 2023; Kerr et al., 2019, 2022; Kumar & Johnston, 2024). Increasing accuracy in cellular computations often requires using more energy; cells optimize this tradeoff to adjust effectively to environmental changes (Lan et al., 2012; Mehta & Schwab, 2012). In some cases, the energy involved in this information processing may be a negligible part of the overall organism’s energy budget. In others, the energetic costs of sensing environments and reprogramming behaviour may be considerable. For single-celled organisms in a highly competitive and resource-limited ecosystem, for example, the sensing machinery and resultant regulatory reprogramming may incur substantial energetic costs. Pursuing this example, bacteria engage in energetically costly behaviors, such as active chemotaxis to seek out more favorable surroundings when resources are abundant (Mitchell & Kogure, 2006; Porter et al., 2011). In contrast, when energy resources are limited, bacteria adopt a more conservative strategy, prioritizing the fundamentals of cellular maintenance over energy-expensive activities (Iuchi & Lin, 1993). In animals, when sensing costs are minimal, informed decision-making via precise foraging or predator avoidance is advantageous, as highlighted by (Lima & Dill, 1990). The balancing act between information-seeking and energy conservation is a fundamental aspect in understanding behavior within dynamic ecosystems from bacterial bet-hedging (Braetz et al., 2017; Shan et al., 2017; Veening et al., 2008) to animal behaviour (Boyles et al., 2020; Humphries et al., 2003; Pyke, 1984).

Organisms from bacteria to eukaryotes are equipped with machinery that can sense and respond to environmental conditions, and changes, in a variety of ways. In quantifying these mechanisms, biomathematics often invokes the picture of proportional-integral-derivative (PID) control, first described mathematically by Nicolas Minorsky in 1922 (Minorsky., 1922). PID control involves sensing and responding to a combination of current state (P), integrated history over time (I), and current rate of change (D). One hundred years later, researchers in synthetic biology are exploring, and engineering, PID control systems within living cells (Aoki et al., 2019; Briat et al., 2016, 2018; Chevalier et al., 2019; Frei et al., 2022). Control systems are naturally employed at the molecular level within living organisms; famous examples include integral control in chemotaxis in *E. coli* (Barkai & Leibler, 1997; Yi et al., 2000), and derivative control of tumbling frequency in bacteria (Alon et al., 1998). PID control offers an effective approach for handling real-world control challenges (Borase et al., 2021), including the noisy environment of the cell (Modi et al., 2021). However, as with other examples of decision-making regulatory machinery in the cell, this capacity comes at an energetic cost, and the availability of energy itself influences decision-making capacity (Forrest et al., 2023; Kerr et al., 2019, 2022; Kumar & Johnston, 2024).

An entire field of work has shed tremendous light on how populations of organisms interact with their dynamic environments (Chesson & Huntly, 1997; Levin, 1976; Levins, 1968; López-Maury et al., 2008), including bet-hedging (Kussell & Leibler, 2005) by switching between phenotypic states (Veening et al., 2008). Here, using a PID picture, we consider the mechanisms by which simple information processing and feedback control can manifest organismal strategies in different environments. We model organisms with two primary behaviors: a resting state and active environmental exploitation. The term ‘resting state’ refers to a state in which an organism reduces its activity and conserves energy (persister states, dormancy, and hibernation fall into this class). In contrast, ‘active environmental exploitation’ is a condition in which organisms actively interact with their surroundings to gain resources. Our model explicitly places the balance between these states under feedback control, similarly to a recent model picturing this balance as analogous to a financial bet-hedging picture (Browning et al., 2019). We explore which forms of feedback control are most beneficial for specific and general environmental behaviours, and how the benefits of sensing and processing information trade off against the energetic costs of this processing.

## Model

Our coarse-grained model is a discrete-space, continuous-time Markov chain with transition rates that depend on environment and potentially on an organism’s particular strategy for environment sensing and feedback control (Fig. 1). The system has an inactive (*I*) and an active (*A*) state, as in the structure of the bet-hedging model in (Browning et al., 2019).

**Fig. 1.**
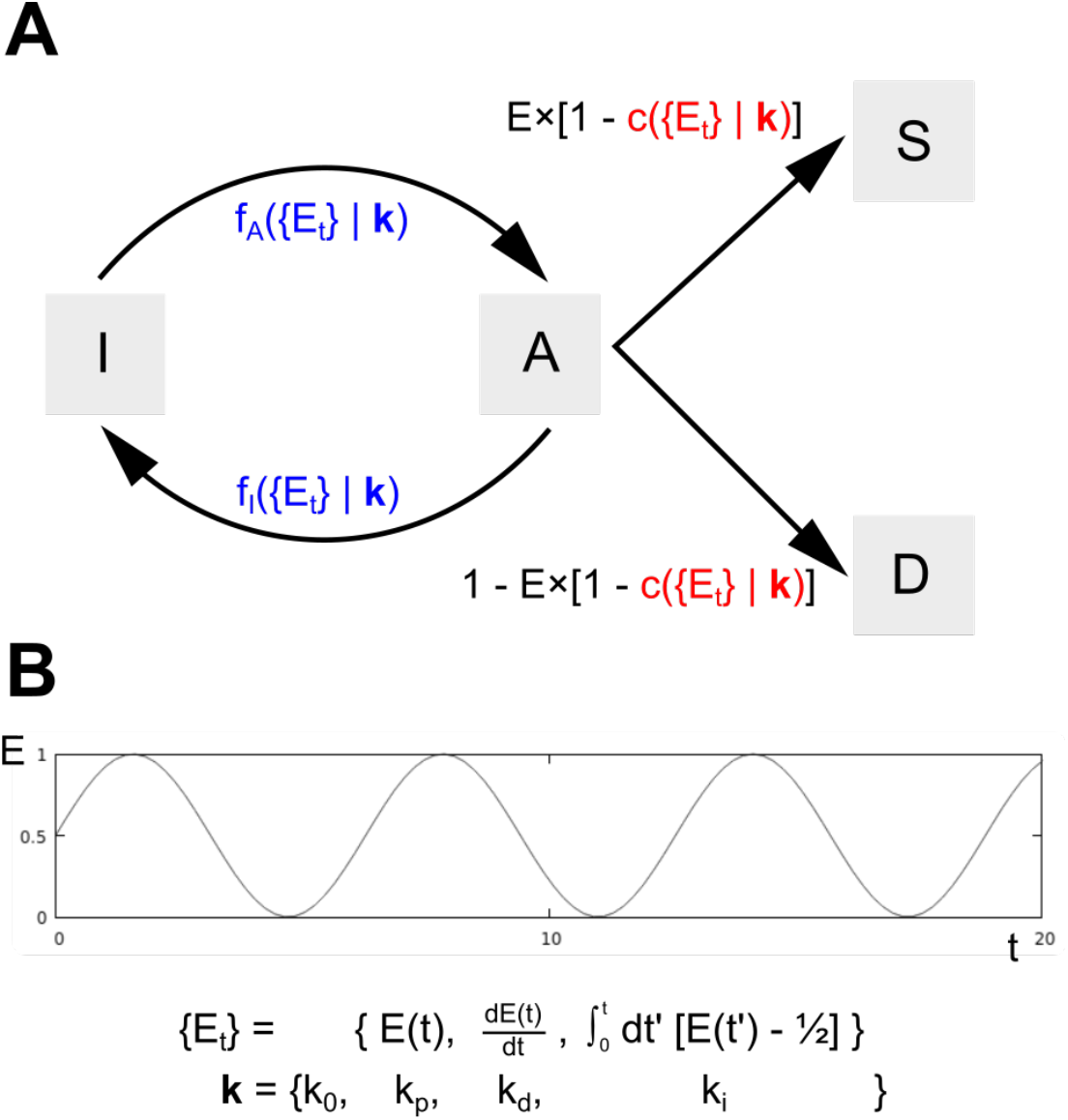
System dynamics. **(A)** State space and dynamics of the system. States are Inactive, Active, Survived, and Died. Transition rates between *I* and *A* (here written *fA* and *fI*) are potentially controlled in response to a collection of environmental behaviour {*Et*} sensed at time *t*, depending on control parameters **k**. Transitions from *A* to *S* are a combination of (positive) current environmental state *E* and (negative) any cost *c* that is incurred through the implementation of control; flux from *A* that does not go to *S* goes instead to *D*. Any negative flux values that arise are set to zero. **(B)** Environmental behaviour and control parameters. An example environmental time series *E*(*t*), bounded on the unit interval and centred on *E* = ½.The collection of environmental behaviour {*E*t} that can be sensed at time *t* consists of the instantaneous value, the instantaneous derivative, and the integral since *t* = 0 of the deviation from *E* = ½. The PID control term is these terms scaled by constants *kp, kd, ki* respectively. *fA* is the control term added to *k0*; *fI* is the control term subtracted from *k0*.

Inactive cells (or organisms) are quiescent, still viable but not capable of reproductive success. Examples may be persister cells in bacterial populations or hibernating animals. Active cells (or organisms) may, if the level of environmental resource available is sufficiently high, experience reproductive success (*S*), but may, if environment resource is limited, die (*D*). The picture here is that the Inactive state is a safe position with no chance of success, while the Active state is a gamble. The key question facing an organism is: when should I be active and when should I be inactive, potentially given some information about the environment?

The equations of motion, reflecting the transitions in Fig. 1, are

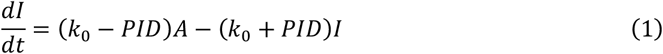

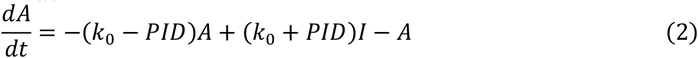

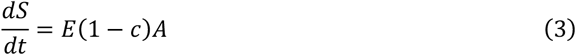

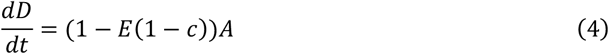

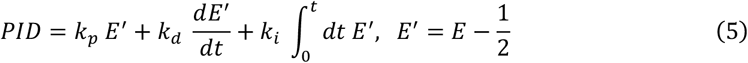

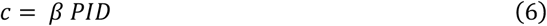

where the cost *c* is an instantaneously-paid penalty, incurred through PID control as the system occupies the active state. The parameter *β* sets the scale of this cost. The environmental signal *E(t)* represents the amount of resource available from the environment. Eqns. 3-4 enforce that “active” organisms *A* experience success *S* with a rate proportional to resource availability minus any incurred energetic cost of sensing; unsuccessful active organisms instead enter the death state *D*. Eqns. 1-2 enforce that the balance between inactive state *I* and active state *A* can be controlled by a PID term sensing the different aspects of environmental dynamics. The environmental signal generally is bounded on the unit interval and has average ½, and when simulated in discrete timesteps takes the form

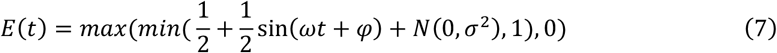

In other words, the deterministic part of Eqn. 7 is computed at every discrete timestep and then, for the stochastic case, an independent and identically distributed normal random variate is added to that value at every timestep. The signal is constrained to lie on the unit interval.

Our null model has the *I*-*A* and *A*-*I* transitions symmetric and independent of environment with rate *k*_0_. Alternatively, organisms can use a form of proportion-integral-derivative (PID) control to help this decision based on their environment (Fig. 1; Eqn. 5). The *I*-*A* transition rate can then be influenced by current environment (proportional), the rate and direction of change of the environment (derivative), and the total historical departure from the average (integral). We use a simple forward Euler method to solve the system in this case. The code for simulation and visualization is freely available at https://github.com/StochasticBiology/energy-sensing.

## Results

### Behaviour of the system

We first show some illustrative behaviours of the system under different conditions. In Fig. 2A, the null case of zero sensing, symmetric *A*-*I* transitions, and constant environment is shown: here the proportion of successful organisms is simply the amount of environmental resource available. In Fig. 2B, the same organisms lacking feedback are exposed to an oscillating environmental resource.

**Fig. 2.**
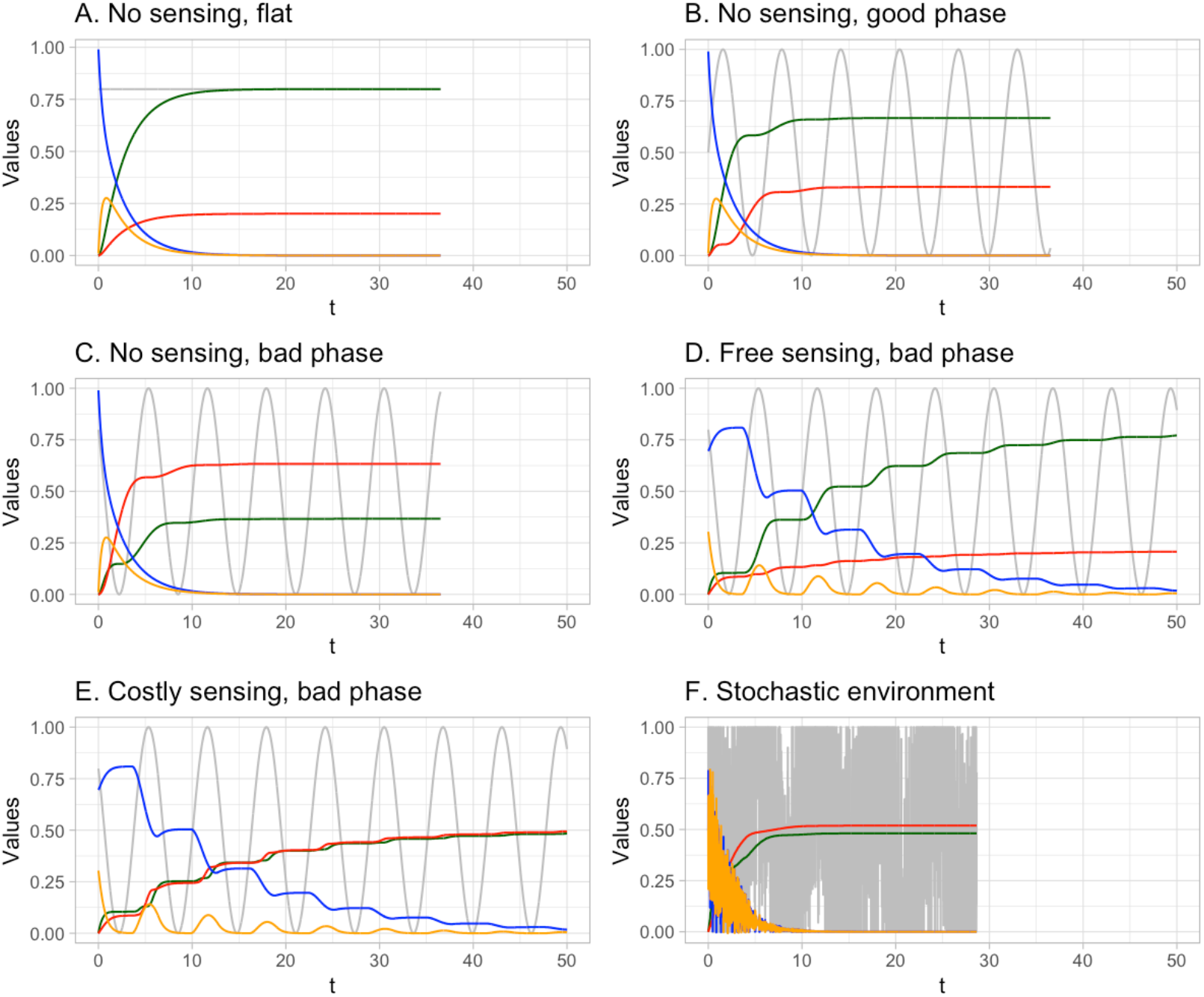
Example time series. Further description in the main text. Traces show population in Inactive (blue), Active (orange), Success (green) and Death (red) states, along with environments resource (grey). **(A)** Constant environment, no sensing. **(B)** Oscillating environment, no sensing. **(C)** Oscillating environment without beneficial phase, no sensing. **(D)** Oscillating environment without beneficial phase, free sensing. **(E)** Oscillating environment without beneficial phase, costly sensing. **(F)** Stochastic environment.

The *A*-*S* and *A*-*D* transitions occur more respectively when resource is below and above average, and the long-term behaviour is reached. In Fig. 2C, exactly the same setup is shown, but for a different environmental resource phase. Now the early evolution of the system takes place in a time of limited resource, and substantial death has occurred before conditions become more favourable. The long-term limit here has only a minority of successes. In Fig. 2D, the same conditions are faced by organisms capable of sensing and controlled their *I*-*A* transitions in response to the environment. Now, *I*-*A* transitions only occur when resource is high, so death is limited, and the long-term behaviour has a large majority of success. In Fig. 2E, exactly the same system is shown but with an energetic cost associated with sensing and responding to the environment. Now the *I*-*A* transitions occur at judicious times, but there are more *A*-*D* transitions because of the incurred resource cost of the sensing and control. Fig. 2F gives an example of the system’s behaviour in a stochastic environment, where the core oscillations are subject to substantial fluctuations.

### Closed-form solution and behaviour of null model

By solving the eigenvalue problem for the equations of motion for *I* and *A*, then using the *A* solution with the sinusoidal driving term from the environment to solve for *S*, an exact solution can be found for the case without feedback. The general solution is too lengthy to be of intuitive value but the long-term behaviour in the representative case of *k*_0_ = 1 is given by

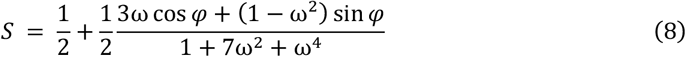

Fig. 3 shows the long term behaviour from Eqn. 8 and from numerical simulation. Intuitively, the most challenging case is low-frequency environmental variation with inopportune phase, so that the system spends a long time with a challenging environment. At the zero frequency limit, Eqn. 8 simply recovers the environmental time series *S* = *E* = ½ + ½ sin φ. As frequency increases, phase dependence is reduced, as the system experiences a more time-averaged picture as the 1/*ω*^4^ term dominates and *S* behaviour tends towards ½. Fig. 3 shows the interdependence of frequency in phase in the optimal (and least optimal) performances, visible in the terms coupling the two variables in Eqn. 8 and reflecting the integrated time of the system spent in a low-resource environment.

**Fig. 3.**
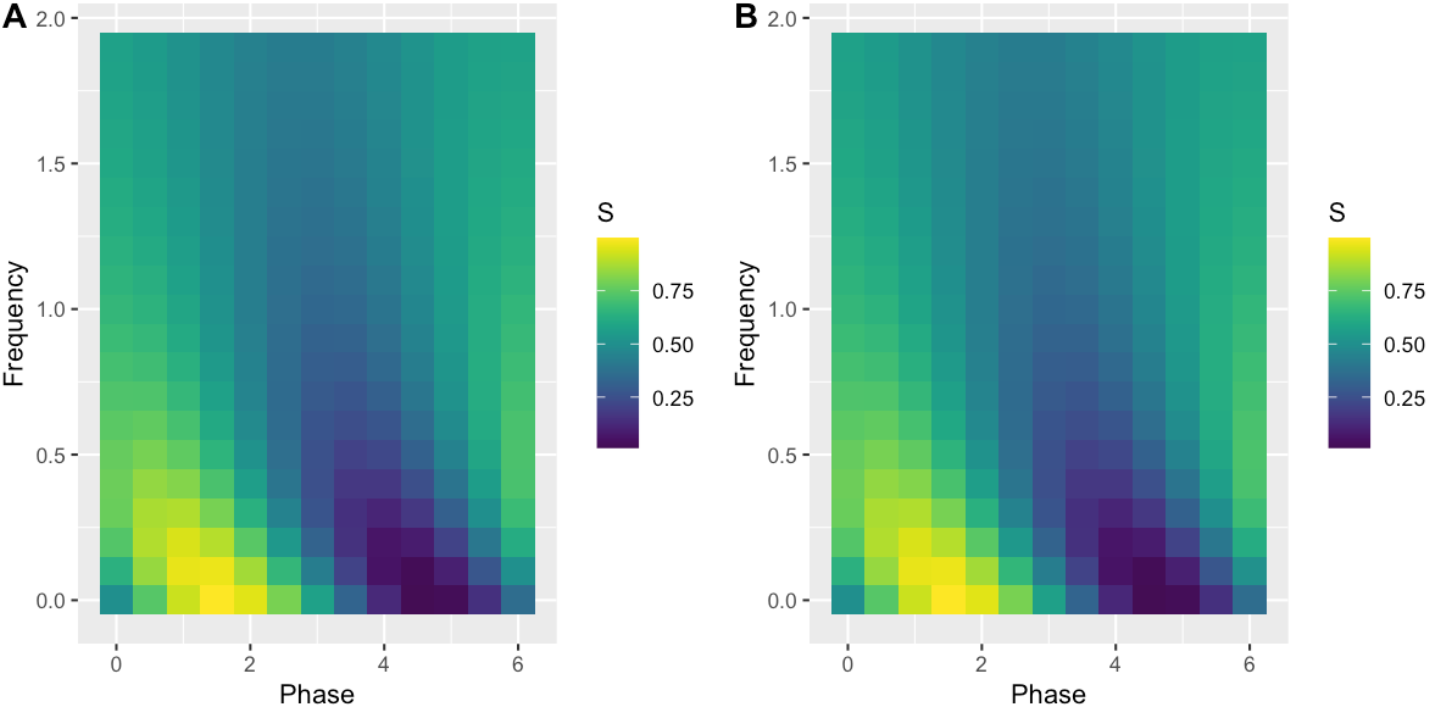
Simulation and analytic results for performance without sensing. Success proportion *S* in the long-term behaviour of the system without sensing, as a function of environmental frequency and phase. (A) Simulation; (B) Eqn. 8.

### Locking-in to the inactive state

We next asked how the system behaved across a range of parameterisations of the PID control model. We scanned through values of *k*_0_, *k*_*p*_, *k*_*i*_, *k*_*d*_ for each environment and recorded (a) whether the system reached a long-term state where all mass was in the *S* and *D* states, and (b) the mass in the *S* state when such a long-term state was reached.

The systems that did not reach a long-term state containing only *S* and *D* exhibited “locked-in” behaviour, with nonzero occupancy of the I state and zero transition rate from *I* to *A* (Fig. 4). Two trivial reasons caused some of these instances: either a collection of zero parameters (including *k*_0_), or a fixed environmental resource under ½ and a collection of parameters (for example, *k*_*p*_, *k*_*d*_) that require higher values for a nonzero transition rate.

**Fig. 4.**
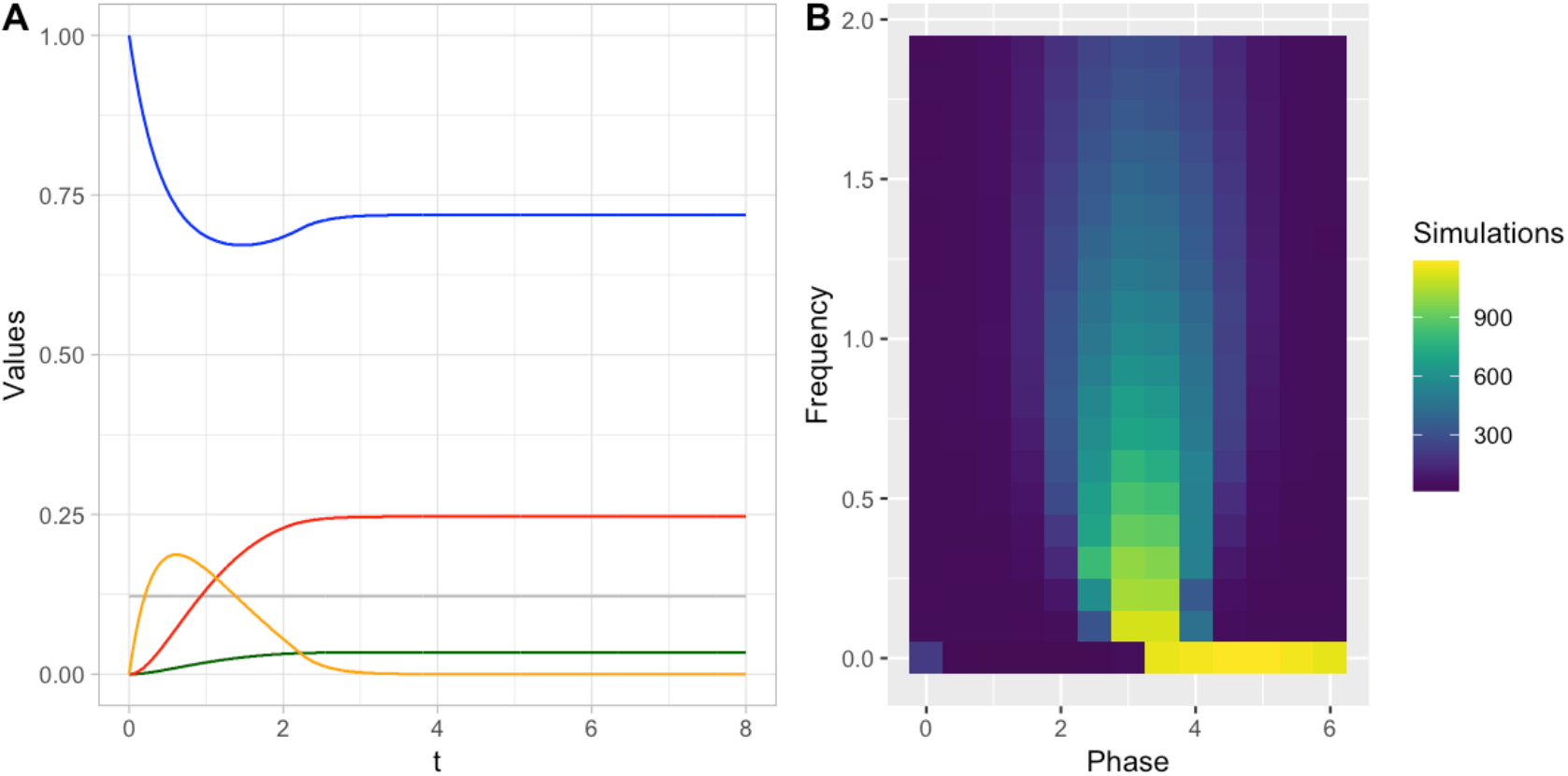
“Locked-in” parameterisations with a nonzero occupancy of the *I* state in the long term limit. **(A)** Example time series as in Fig. 2; traces show population in Inactive (blue), Active (orange), Success (green) and Death (red) states, along with environmental resource (grey). In this example *β* = 0; *ω* = 0; *φ* = 4.0; *k0* = 1; *kp* = 0.4; *ki* = 1; *kd* = 0.4. (**B)** Number of simulations with nonzero long-term occupancy of the *I* state with environmental phase and frequency.

The less trivial circumstance leading to this “locked-in” behaviour involved integral control. Here, the other aspects of the controller would support some transitions to the *A* state. But the integral control term, increasing in magnitude over time, grows to overcome the contributions of the other terms. Hence, faced with a long period of below-average resource, the resource commits completely to the inactive state. An equally long period of above-average resource would be needed to bring the integral control term back to zero (or positive) values and support more transitions to the active state – hence, the longer the low-resource period, the more high-resource time is required to risk activity.

### System behaviour with free or costly sensing

Of the parameterisations that did give long-term behaviour with zero *I* and *A* content, we next asked what the optimal control strategy was for a given environment. Fig. 5 shows the optimal performance (over *k*_0_, *k*_*p*_, *k*_*i*_, *k*_*d*_) that we found for a given environmental resource behaviour, and the scale of improvement over the null case without sensing in Fig. 3. Sensing (without cost) allows a substantial improvement across many environments, particularly those with phases that challenge the null case (as in Fig. 2). The advantages are most pronounced at lower frequencies, but even at higher frequencies there exist parameterisations that give a substantial performance increase. In the case of costly sensing, the picture remains broadly the same, but with lower peak performance (because the cost of sensing reduces success probability).

**Fig. 5.**
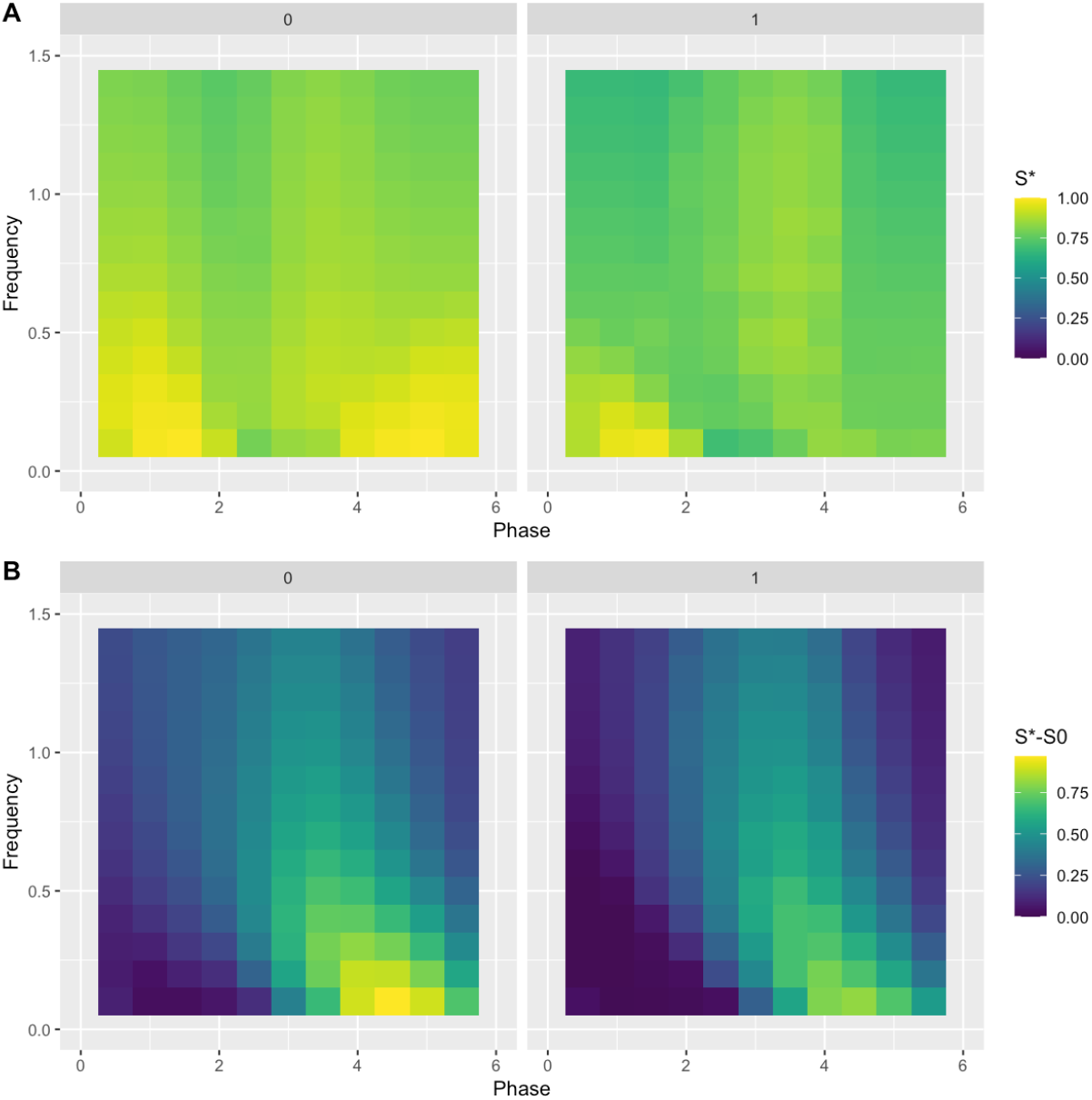
Performance with feedback control. **(A)** Optimal performance *S\** with sensing, for (left) free sensing *β* = 0 and (right) costly sensing *β* = 1. **(B)** Improvement over case *S0* without sensing (illustrated in Fig. 3).

We next asked what were the specific control parameters that gave rise to these optimal behaviours. The contributions of baseline (open-loop) control and the different aspects of the PID controller were rather complex (Fig. 6).

**Fig 6.**
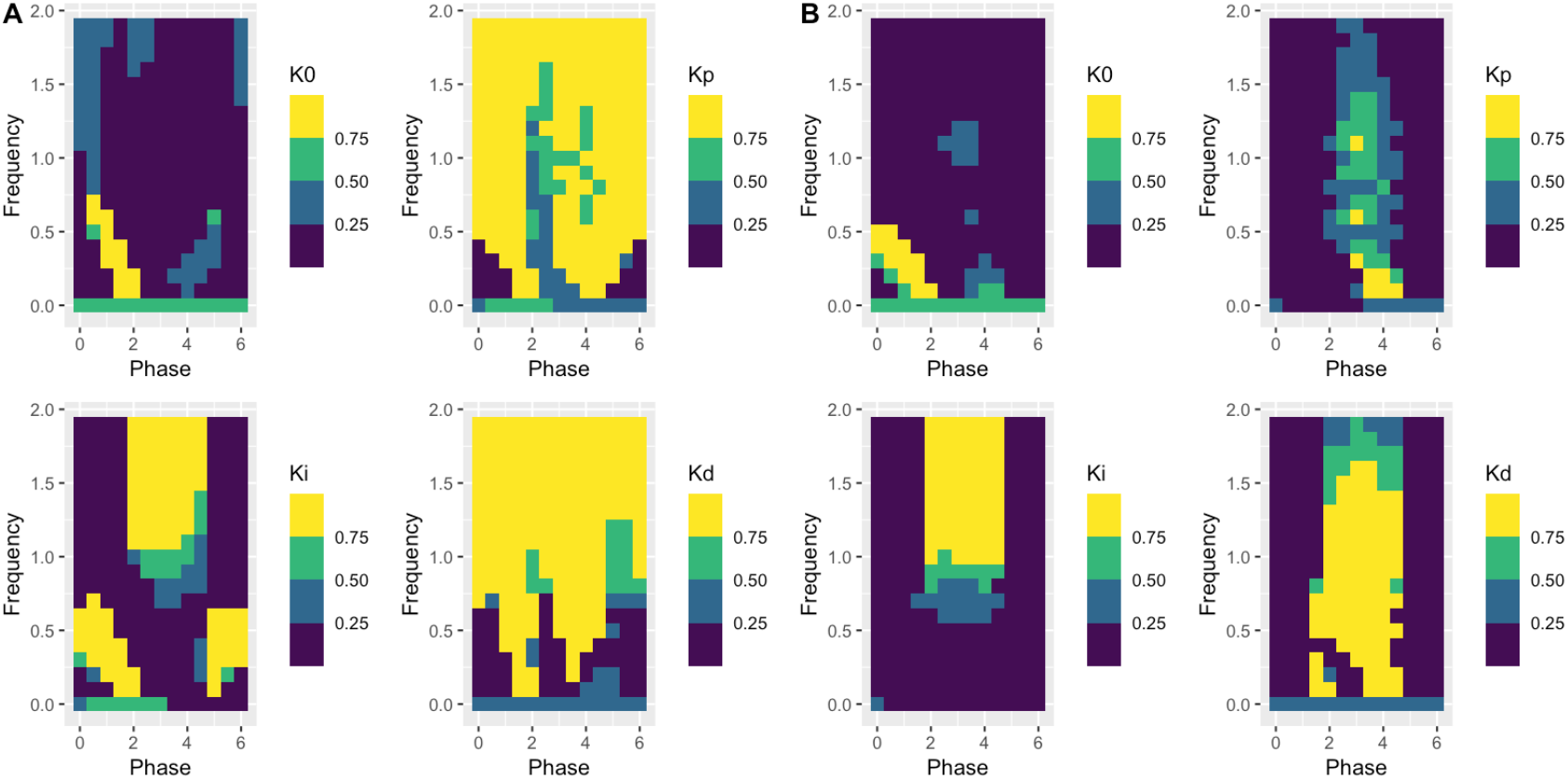
Optimal strategy of *k*_0_, *k*_*p*_, *k*_*i*_, *k*_*d*_ for a given environment. (A) free sensing *β* = 0 and (B) costly sensing *β* = 1.

In almost all cases with zero sensing cost, proportional and derivative controls take maximal values. The exception is for low-frequency cases with phases so that resource starts with a high value. Here, no control terms are assigned a particularly high value, as there is little advantage to any strategy other than adopting the active state as early as possible. The integral term also increases for high-frequency cases where the phase starts with low resource. The optimal baseline term is low except in the case of zero frequency or the best possible environmental situation – in these cases there is respectively no advantage to sensing (absent) changes, and no particular advantage to specific sensing mechanisms.

When sensing is costly, these patterns of optimal parameterisations shift somewhat. Proportional and derivative control is retained only for phases where resource starts low. The integral control component is absent at low frequencies, appearing only at higher frequencies for low-starting phases. The open-loop term (which does not correspond to sensing, and hence does not contribute to cost) is relatively high across all cases.

These results demonstrate the specific best control strategy for a given environment. We next asked which control strategies had the best performance across the whole range of environments we consider. In other words, which strategies are good regardless of which environment the system faces?

Fig. 7 shows the results. In the zero-cost case, the favouring of proportional and derivative control observed across Fig. 6 is clear. In the costly case, the best strategies also involve proportional and derivative control. Integral control is less apparent in the costly case, as it can in principle incur a greater cost: it accumulates information over time and therefore accumulates cost over time as well. The merit of integral control would seem to be clearest in the “locking-in” behaviour shown previously, where it supports long-term inactivity in response to long periods of suboptimal resource availability.

**Fig. 7.**
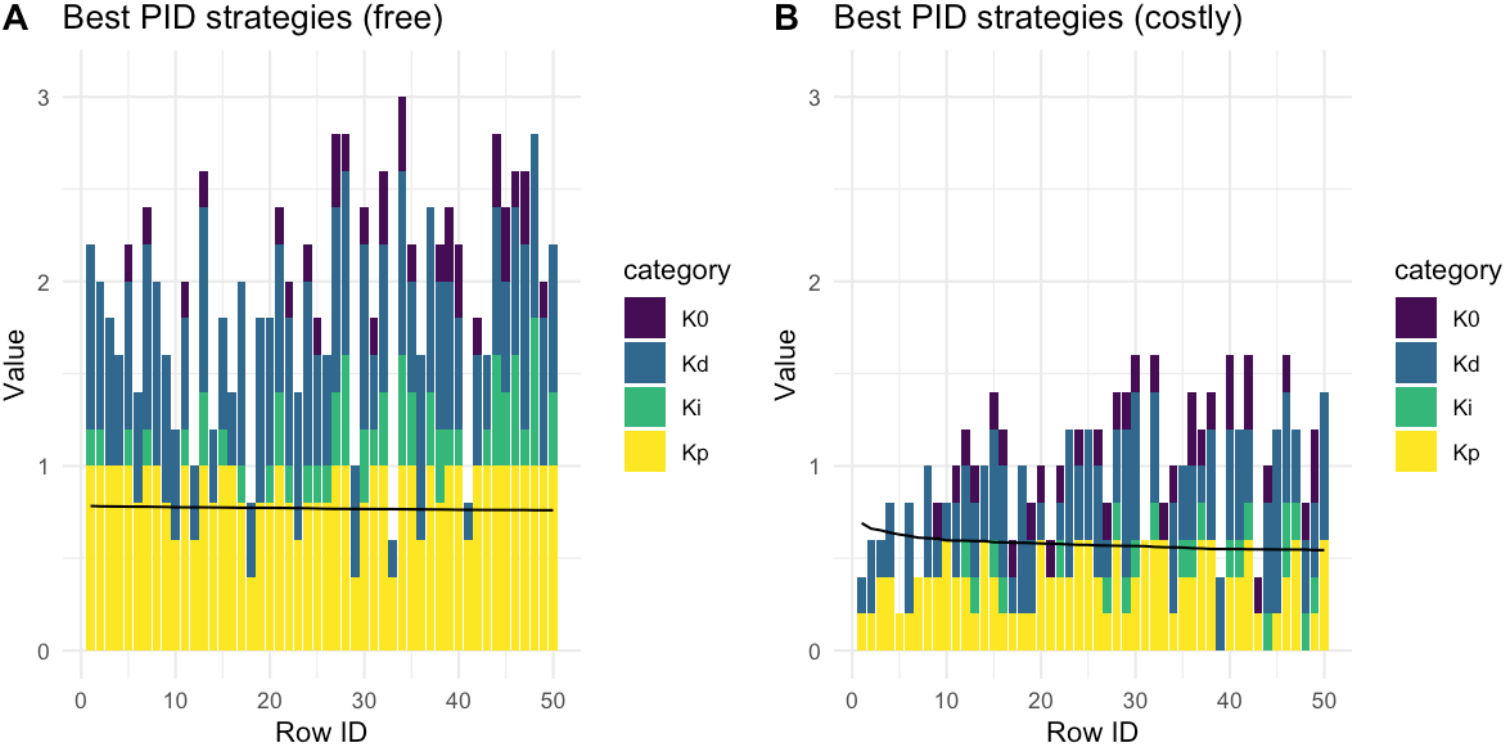
Generally good control strategies. Optimal strategy across all environments, visualized as stacked columns describing the base and PID parameterization for each strategy, with **(A)** free sensing *β* = 0 and **(B)** costly sensing *β* = 1. The height of each coloured bar is that parameter’s value in that parameterization. “Row ID” simply ranks strategies by their average performance across all environments, given by the black line.

### System behaviour in stochastic environments

We next asked how the system performed in stochastic environments, capturing this by layering fluctuations of a given magnitude additively over the time series investigated previously. Generally, performance is lower in stochastic cases, but sensing still allows a substantial and general improvement over the null case that has the same shape as the deterministic case and is limited by costly sensing (Fig. 8, Supp. Fig. 1).

**Fig. 8.**
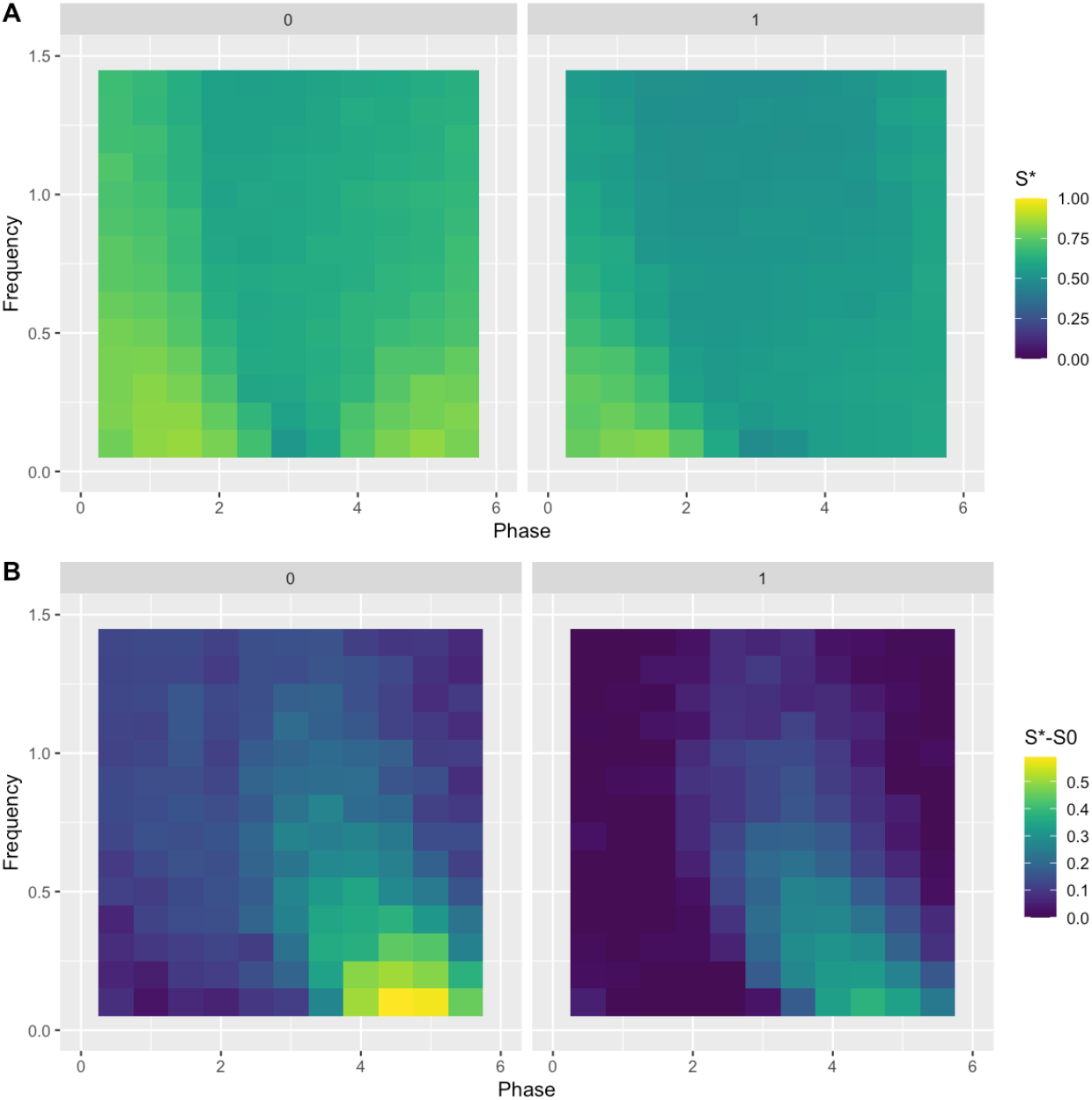
Performance with sensing in the stochastic system. Here the scale of environmental noise ε = 0.5, as in Fig. 2F. **(A)** Optimal performance *S\** with sensing, for (left) free sensing *β* = 0 and (right) costly sensing *β* = 1. **(B)** Improvement over case *S0* without sensing (Fig. 3). Similar behaviours, scaled intuitively, are observed for different values of cost and noise magnitude (Supp. Fig. 1).

A notable difference when exploring the optimal feedback parameters for the stochastic case is the (intuitive) disappearance of the derivative control. Because the noise is uncorrelated, the derivative term contains less useful information about the current and future behaviour of environmental resource. When this uncorrelated noise dominates the time behaviour, the value of the derivative term is negligible. The proportional term is clearly present across the free-sensing cases. An integral term is also present, allowing the system to remain inactive when the nutrient resource remains below average for a time period. However, the advantage that this sensing provides is relatively marginal, and average performance across environmental situations is not very different from the null case (Fig. 9, Supp. Fig. 2).

**Fig. 9.**
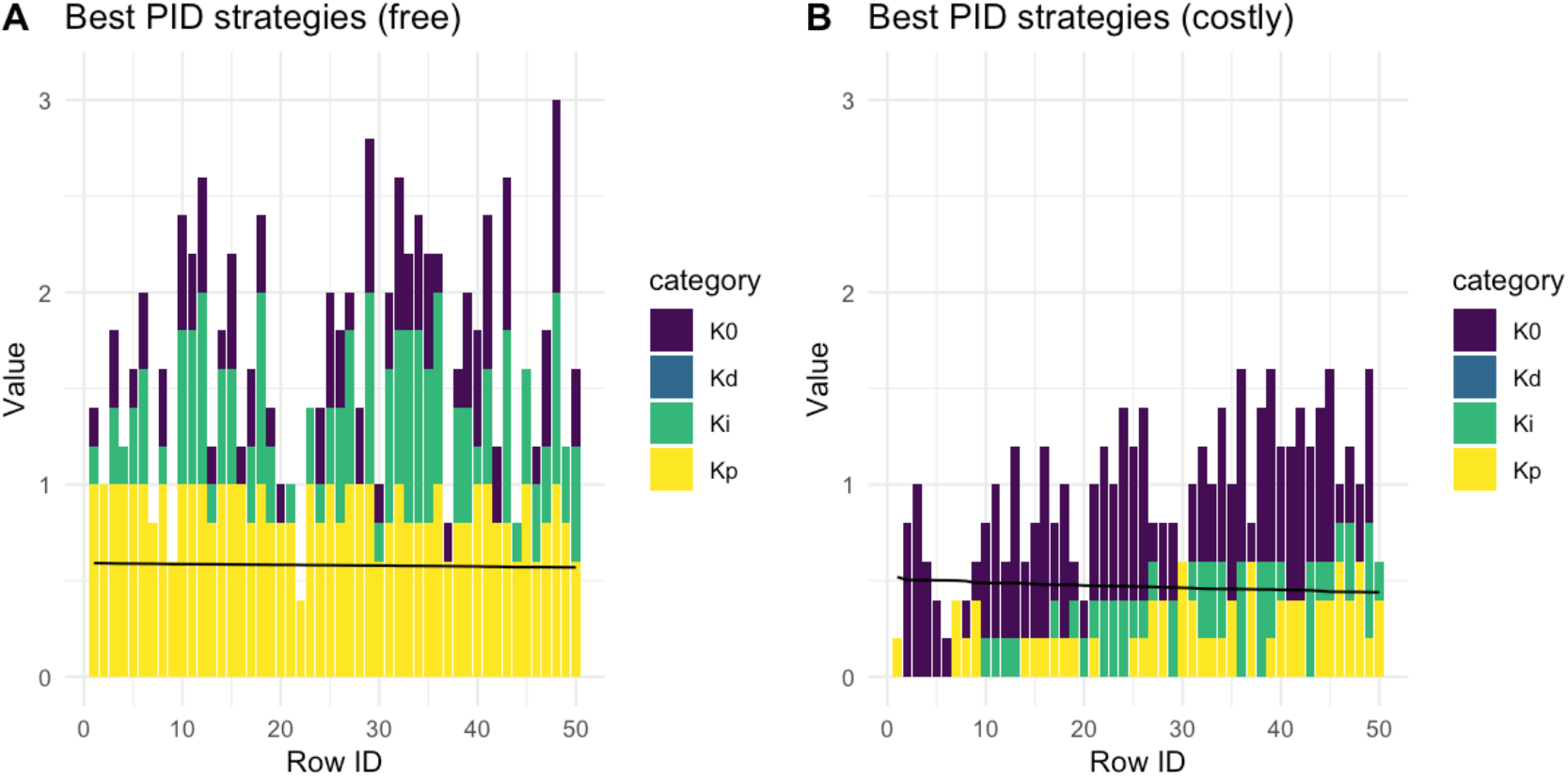
Generally good control strategies for the stochastic system. Optimal strategy across all environments, with **(A)** free sensing *β* = 0 and **(B)** costly sensing *β* = 1. The height of each coloured bar is that parameter’s value in that parameterization. “Row ID” simply ranks strategies by their average performance across all environments, given by the black line. Similar behaviours, scaled intuitively, are observed for different values of cost and noise magnitude (Supp. Fig. 2).

Accordingly, only weak to zero parameterisations of the PID controller are preserved in the costly sensing case (Fig. 9, Supp. Fig. 2).

## Discussion

Energy is required for cellular decision-making. Previous work at a finer-grained scale has demonstrated the importance of ATP supply in cellular decision-making circuitry (Forrest et al., 2023; Kerr et al., 2019, 2022; Kumar & Johnston, 2024). That work has shown that without sufficient energy supply, cellular decision-making capacity is more limited. This work attempts to link with a coarser-grained picture. Given that energy is required to sense, process, and respond to an environment, under what circumstances is this price worth the reward?

The parameterization of our model is deliberately left quite general, in order to demonstrate the general properties of a system that may or may not use sensing and feedback to change its “strategy” in the face of changing environments. Quantitative connections with real biological systems must fix a particular interpretation of resource and risk. For example, for bacterial cells, resource could be nutrient concentration in the environment, and risk could be outcompetition from rivals in a growing population (Tufto, 2015; Veening et al., 2008). Then, the costs of sensing and processing information (and responding, perhaps by changing a large-scale transcription programme) may be a substantial part of the overall cellular energy budget. For animals, resource could be available nutrition, with the associated risk of insufficient energy to survive (Boyles et al., 2020; Humphries et al., 2003). Hibernation is an clear case of inactivity over a (long) period of limited resource availability; diurnal patterns of dormancy (depending on the source of nutrition) are an example on a shorter timescale.

What general principles can we distil from the specific picture presented here? First, even if environmental sensing incurs costs on the scale of resource availability, feedback control can still provide substantial benefit to organisms. Integral control can be employed to “lock in” a state of inactivity when faced with an extended period of low resource (Fig. 4), to lock in a state of activity in an extended period of high resource, or to average over short-time fluctuations in a stochastic environment (Fig. 9A). Proportional sensing, intuitively, provides an immediate response to a given environment, and is typically present in most parameterisations that perform well across different environments. In the absence of sensing cost, proportional control is almost universally adopted across environments (Fig. 6A). In predictable environments, differential control is also typically adopted (Figs. 6-7), allowing the system some ability to predict the future behaviour of the environment. In stochastic environments, differential control is abandoned, as the instantaneous change in resource availability is not correlated with future behaviour and therefore does not contain useful information (Fig. 9). The extent to which different *costly* control strategies are favoured depends on the particular situation. Strikingly, even if the resource invested in sensing is not dissimilar in scale to the total resource availability, nonzero control strategies can still be favoured in several circumstances (Figs. 6-9). However, the improvement from sensing averaged across different environmental conditions is relatively small in the stochastic case compared to the deterministic case (Fig. 9, Supp. Fig. 2).

A tighter connection to specific biological systems will of course be desirable to take this work forward. It is not impossible that the costs of sensing and responding to the environment (for example, activating different transcriptional programmes) could be quantified, or at least estimated (Johnston et al., 2014; Phillips & Milo, 2009), experimentally in simple systems like bacteria. Parameters from such experiments would allow this modelling approach to be adapted to a specific biological system, and its predictions about adaptable strategies to be tested in further experimental work.

## Supplementary Information

**Supplementary Figure 1.**
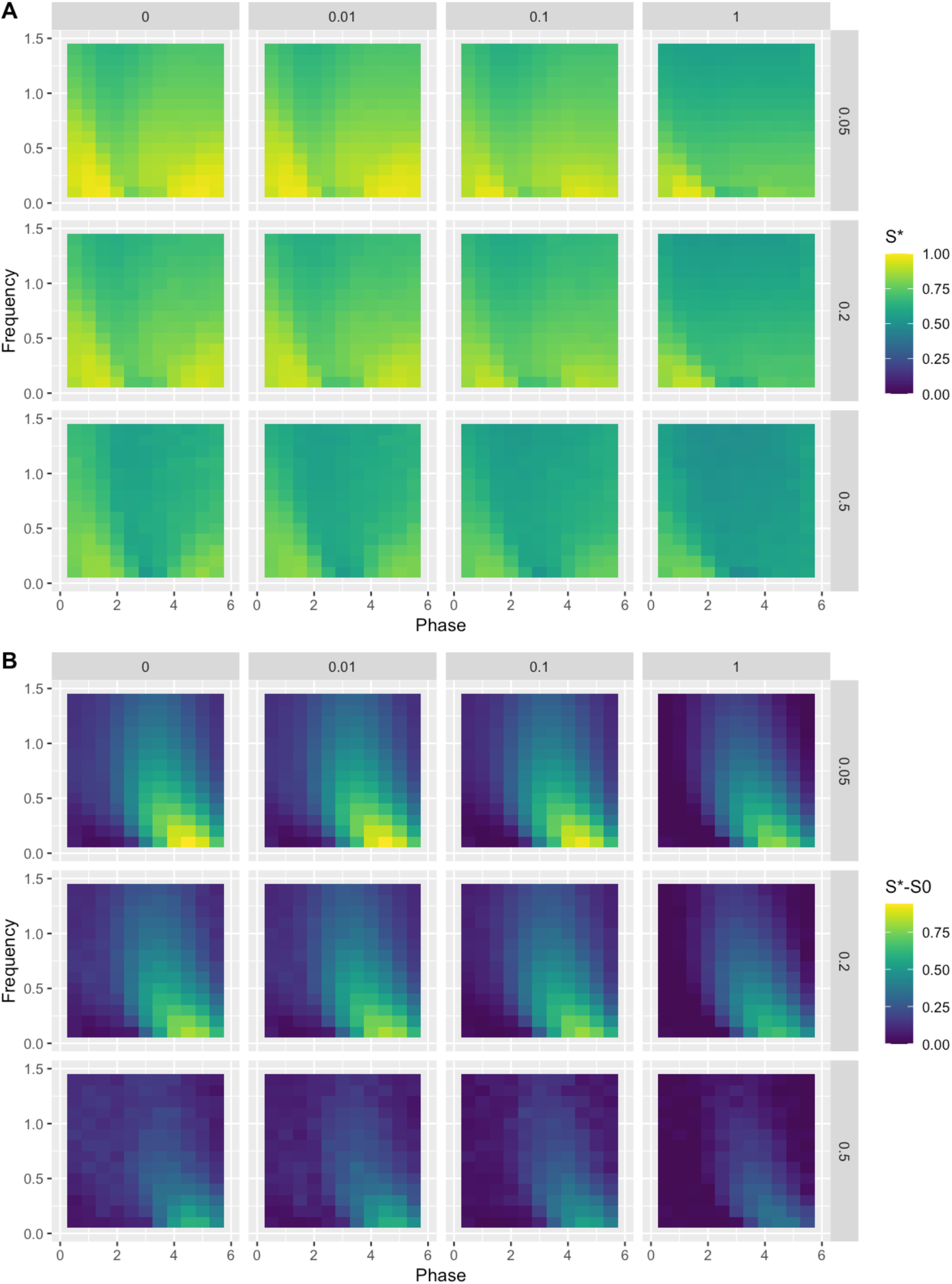
Optimal performance of the system for a wider variety of cost scales β (columns) and scales of noise ε (rows).

**Supplementary Figure 2.**
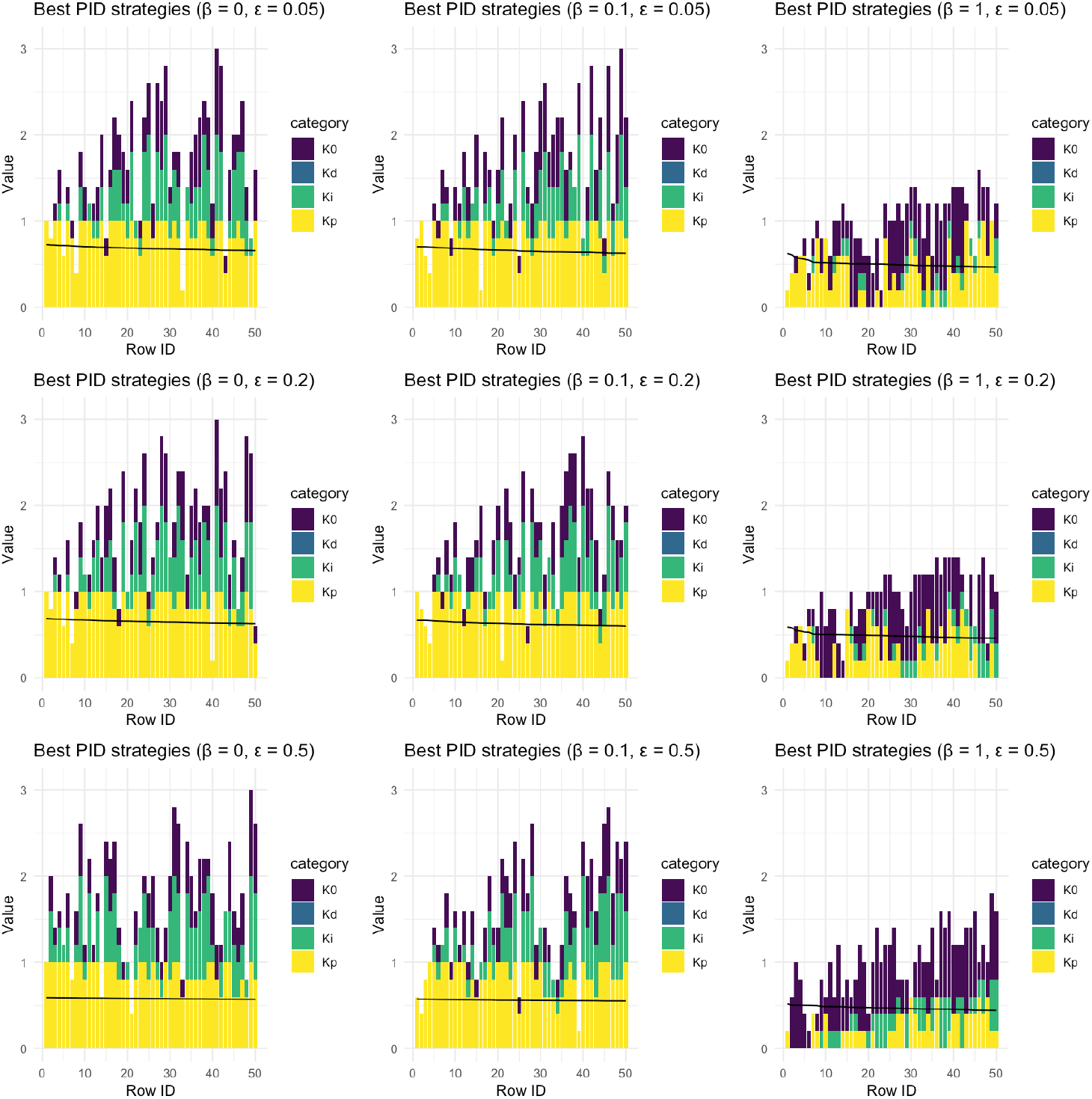
Optimal PID parameterisations across environmental behaviours, for a wider variety of cost scales β (columns) and scales of noise ε (rows).

